# FaStore – a space-saving solution for raw sequencing data

**DOI:** 10.1101/168096

**Authors:** Łukasz Roguski, Idoia Ochoa, Mikel Hernaez, Sebastian Deorowicz

## Abstract

The affordability of DNA sequencing has led to the generation of unprecedented volumes of raw sequencing data. These data must be stored, processed, and transmitted, which poses significant challenges. To facilitate this effort, we introduce FaStore, a specialized compressor for FASTQ files. The proposed algorithm does not use any reference sequences for compression, and permits the user to choose from several lossy modes to improve the overall compression ratio, depending on the specific needs. We demonstrate through extensive simulations that FaStore achieves a significant improvement in compression ratio with respect to previously proposed algorithms for this task. In addition, we perform an analysis on the effect that the different lossy modes have on variant calling, the most widely used application for clinical decision making, especially important in the era of precision medicine. We show that lossy compression can offer significant compression gains, while preserving the essential genomic information and without affecting the variant calling performance.

## Introduction

The growing interest in applications of genome sequencing, together with the dropping costs and continuous improvements in sequencing technologies, has led to the generation of unprecedented volumes of increasingly large and ubiquitous raw genomic data sets (Stephens et al. 2015). These data are characterized by highly-distributed acquisition, massive storage requirements, and large distribution bandwidth. For example, the 1000 Genomes Project (Clark et al. 2012) required 260 Terabytes of storage space (for raw, aligned, and variant calling data). Moreover, the 100,000 Genomes Project (2017) has already exceeded 21 Petabytes in size. Such flood of data hampers the efficiency of data analysis protocols, limits efficient data sharing, and generates vast costs for data storage and IT infrastructure (Schadt et al. 2010). This situation calls for state-of-the-art, efficient compressed representations of the raw genomic data, that can not only alleviate the storage requirements, but also facilitate the exchange and dissemination of these data.

The raw high-throughput sequencing data are primarily stored in FASTQ files (Cock et al. 2010), which are usually considered as the input for the genomic data processing and analysis pipelines. A FASTQ file can be perceived as a collection of “reads”, each containing a sequence of nucleotides (generally referred to as the read), the quality score sequence that indicates the reliability of each base in a read, and the identifier, which usually contains the information about the sequencing instrument, flow cell coordinates, etc. Millions of such reads are produced in a single sequencing run. As a result, storing raw whole-genome sequencing data of a single human can easily exceed 200 GB in uncompressed form.

Most of the analyses pertaining to DNA sequencing (e.g., in the context of precision medicine) rely on assessing the variants of the sequenced genome against a known reference genome. In order to assess these variants, the raw reads present in the FASTQ file are first aligned to a reference sequence. This process generates aligned reads in SAM format (Li et al. 2009), which contains the same information as the FASTQ file (i.e., the read identifier, the raw sequence of nucleotides and the quality scores), together with the alignment information for each read, and possible additional information provided by the mapper. After a number of post-processing steps over the aligned reads, variant calling against the reference genome is performed. The called variants, following a number of additional data cleaning and validation steps, is normally used as an entry point for further clinical analyses. The size of the resulting file is orders of magnitude smaller than the input raw reads stored in FASTQ format. For example, as humans share about 99.5% of the same genetic code (Wheeler et al. 2008), the variants of interest will account for less than 1% of all the base pairs.

Although the data contained in the FASTQ file could potentially be recovered from the corresponding SAM file (or its compressed version), there are cases in which this may be impossible. For example: (*i*) the SAM file may not contain the reads that failed to align to the reference genome or those marked as duplicates, (*ii*) some reads in the SAM file can be truncated (hard-clipped), (*iii*) the reference genome used for compression may no longer be accessible during decompression, or (*iv*) there may not be a reference genome at all to generate the corresponding SAM file (e.g., in metagenomics, the different organisms present in the sequenced sample are generally unknown prior to the analysis).

Therefore, we focus on the compression of the information stored in FASTQ format, that is, the raw data containing the nucleotide sequences, the read identifiers, and the quality scores, since they represent a minimal subset of the data required for future reproduction of the analyses performed on the sequenced data. The existing specialized solutions for FASTQ files compression, extensively examined in (Numanagić et al. 2016), obtain significant compression gains over general compression tools such as gzip. However, in practice, gzip is still the *de facto* choice, mainly due to its popularity and stability. It seems that the community has not decided yet that the assets of specialized FASTQ compressors are worth some complications that may appear when moving to a different format of storage. Also, the fact that currently a number of good dedicated compressors are available does not make the right choice simple.

In this paper we propose FaStore, a new compressor for FASTQ files that, among others, can be used for long-term archival of raw genomic information and efficient sharing, especially with limited internet bandwidth. FaStore inherits the assets of our previous attempts in the field, especially DSRC (Deorowicz and Grabowski 2011; Roguski and Deorowicz 2014), ORCOM (Grabowski et al. 2015), and QVZ (Malysa et al. 2015; Hernaez et al. 2016). The proposed compressor offers both lossless and lossy compression modes (the latter only for the quality scores and the identifiers), and does not use any external reference sequences.

We show that FaStore significantly outperforms the existing compressors in the lossless mode. We, however, advocate for the lossy option when suitable, which, as presented, gives much better shrinkage of the input files with negligible differences in variant calling. Although we emphasized the variant calling as the most common use-case for precision medicine, the analyses performed using FASTQ files are not only limited to variant calling, but also used for, e.g., gene expression analysis, assembly, or metagenomics.

## Results

### Lossless and lossy compression of sequencing data

FaStore is a compressor optimized for handling FASTQ files produced by next-generation sequencing platforms, which are characterized by generating massive amounts of short reads with a relatively low sequencing error rate. FaStore exploits the redundancy present in the reads to boost the compression ratio. In addition, it includes several compression modes to account for the different needs that the users may have. In particular, parts of the data, namely, quality scores and read identifiers, can be optionally discarded or quantized for additional file size reduction. On the other hand, the sequences of nucleotides (DNA sequences) are always lossless compressed. Due to the different nature of components of reads (DNA sequences, quality scores, and identifiers), FaStore uses different specialized compression techniques for each of them (see **Supplementary Methods** for details). Moreover, as the sequencing data can be generated from a library in a single- or paired-end configuration, FaStore provides different techniques to handle both cases, guaranteeing that the pairing information between the reads is preserved when available. In the following, when clear from the context, the DNA sequences may be also referred to as reads.

Compression of the DNA sequences is done without the use of any external reference sequences. Relying on a reference sequence for compression requires the availability of the same reference at the time of decompression, which may no longer be accessible, thus making the compressed DNA sequences unrecoverable. Hence, this design choice guarantees perfect reconstruction of the sequences.

The reads produced using next-generation sequencing protocols can be thought of as being randomly sampled from across the genome (Firtina et al. 2016) (the input molecule), and thus their initial ordering in the output file carries no information.

With this in mind, FaStore reorders the reads to exploit the existing high similarity among the DNA sequences. In particular, the reads are clustered in a manner such that reads coming from neighboring positions in the sequenced genome are likely to belong to the same cluster. When possible, within each cluster, the reads are assembled into contigs and stored relatively to a consensus sequence. Alternatively, a read can also be stored relatively to other reads belonging to the same cluster, or “as it is”, depending on the degree of similarity with the other reads in the cluster.

As a trade-off between the computation time to cluster the reads and the attained compression ratio, FaStore offers two modes of operation, denoted by C0 (fast) and C1 (default). The clustering process in C1 mode is more involved, leading to better compression ratios. The decompression speed is similar for both modes.

While the DNA sequences can be efficiently compressed due to the redundancy present in the data, the quality scores have proven more difficult to compress (Bonfield and Mahoney 2013). Part of the reason is that they are inherently noisy and thus characterized by a high entropy. In addition, preserving precise quality scores is often unnecessary (i.e., some distortion is generally acceptable), in that no cost is incurred on the subsequent analyses performed on the data (Ochoa et al. 2016; Yu et al. 2015). Hence, FaStore offers, in addition to lossless compression, various types of lossy compression modes for the quality scores. In particular, FaStore includes Illumina 8-level binning (Illumina 2014), a custom binary thresholding, and an adaptive scheme based on QVZ (Malysa et al. 2015; Hernaez et al. 2016).

Illumina binning mimics the binning of quality scores performed by the latest Illumina sequencing machines by mapping the resolution of quality scores to 8 distinct bins. The binary thresholding quantizes the quality scores according to a user-provided threshold, i.e., it sets the quality values below the threshold to *q*_min_, and those above to *q*_max_. Finally, QVZ quantizes the quality scores so as to minimize the rate allocation (number of bits per quality score) while satisfying a distortion constraint. To design the appropriate quantizers, QVZ relies on computing the statistics of the quality scores prior to compression. FaStore gathers these statistics while clustering the reads, and thus there is almost no added computational cost. The quantizers are generated after clustering the reads, and one global codebook per dataset is used. For lossless compression, FaStore uses QVZ in lossless mode.

The read identifiers are initially tokenized to make use of the fact that some appearing tokens are constant, some are from a small dictionary, etc. Moreover, since the complete identifiers are usually unnecessary in practice, FaStore also offers a lossy mode for storing them, either by removing the comments (as mappers do by default) or by completely skipping them.

Regarding storing of the pairing information between the reads (when these were generated from a paired-end library), the FASTQ format does not clearly define how the pairing information should be represented. Currently there are two main ways to convey this information: (i) it is carried in the read identifiers (i.e., a pair of reads share the same identifier), or (ii) it is encoded at the file-level (i.e., the reads reside on the same lines in two FASTQ files or are stored interleaved in a single file). FaStore preserves this information with the sequences, allowing the identifiers to be removed and generating unique ones per pair of reads when decompressing.

### Compression factors

For evaluation of the proposed compressor FaStore, we use a subset of data sets already benchmarked in (Numanagić et al. 2016; Grabowski et al. 2015; Benoit et al. 2015; Deorowicz and Grabowski 2013), alongside new ones characterized by a high coverage. The details of the employed datasets are summarized in Table 1 (see **Supplementary Methods** for the download links). The collection consists of 7 large sets of pair-end FASTQ files (top 7 rows) and one vast pair-end data set (bottom row), and it includes sequencing data from the *H.sapiens, G.gallus*, and *C.elegans* species. We compared the performance of FaStore with that of gzip (the *de facto* current standard in storage of sequencing data) and the top FASTQ compressors according to (Numanagić et al. 2016): DSRC 2 (Roguski and Deorowicz 2014), Fqzcomp (Bonfield and Mahoney 2013), Leon (Benoit et al. 2015), Quip (Jones et al. 2012), and Scalce (Hach et al. 2012). We also tested the top DNA-only compressors according to (Numanagić et al. 2016): ORCOM (Grabowski et al. 2015), Mince (Patro and Kingsford 2015), and BEETL (Cox et al. 2012). However, since these algorithms fail to compress the whole FASTQ file, we relegate their results to the **Supplementary Worksheet W1**. Unfortunately, for several data sets, Mince run out of available memory (128 GB) and BEETL failed to process some of them in 48 hours time, so they are not included in our analysis.

**Table 1:** Datasets used in the experiments. WGS is for Whole Genome Sequencing. WES is for Whole Exome Sequencing.

| Data set | Species | Experiment type | Sequencer | FASTQ size [GB] | Read len. [bp] | No. reads [M] | Coverage |
| --- | --- | --- | --- | --- | --- | --- | --- |
| CE | *C.elegans* | WGS | Illumina GA II | 17.6 | 100 | 67.6 | 66 |
| GG | *G.gallus* | WGS | Illumina GA II | 115.9 | 100 | 347.3 | 32 |
| HS2 | *H.sapiens* | WGS | Illumina GA II | 336.9 | 100 | 1339.7 | 43 |
| HSX | *H.sapiens* | WGS | Illumina HiSeq X | 292.1 | 151 | 819.1 | 39 |
| WEX | *H.sapiens* | WES | Illumina HiSeq 2500 | 44.7 | 126 | 150.4 | ~220 |
| WGS-14 | *H.sapiens* | WGS | Illumina HiSeq 2000 | 115.6 | 101 | 447.1 | 14 |
| WGS-42 | *H.sapiens* | WGS | Illumina HiSeq 2000 | 337.3 | 101 | 1304.5 | 42 |
| WGS-235 | *H.sapiens* | WGS | Illumina HiSeq 2000 | 1888.4 | 101 | 7363.7 | 235 |

Figures 1a–d show the average compression factor and compression/decompression speeds for the complete collection. All the tested compressors were run using 8 processing threads, when applicable, and in maximum compression mode. Due to space constraints and ease of exposition, we provide results for the main lossy settings of FaStore (denoted by *reduced, lossy*, and *max*), and refer the reader to the **Supplementary Worksheet W1** for an extensive evaluation of the whole range of lossy modes provided by FaStore.

**Figure 1:**
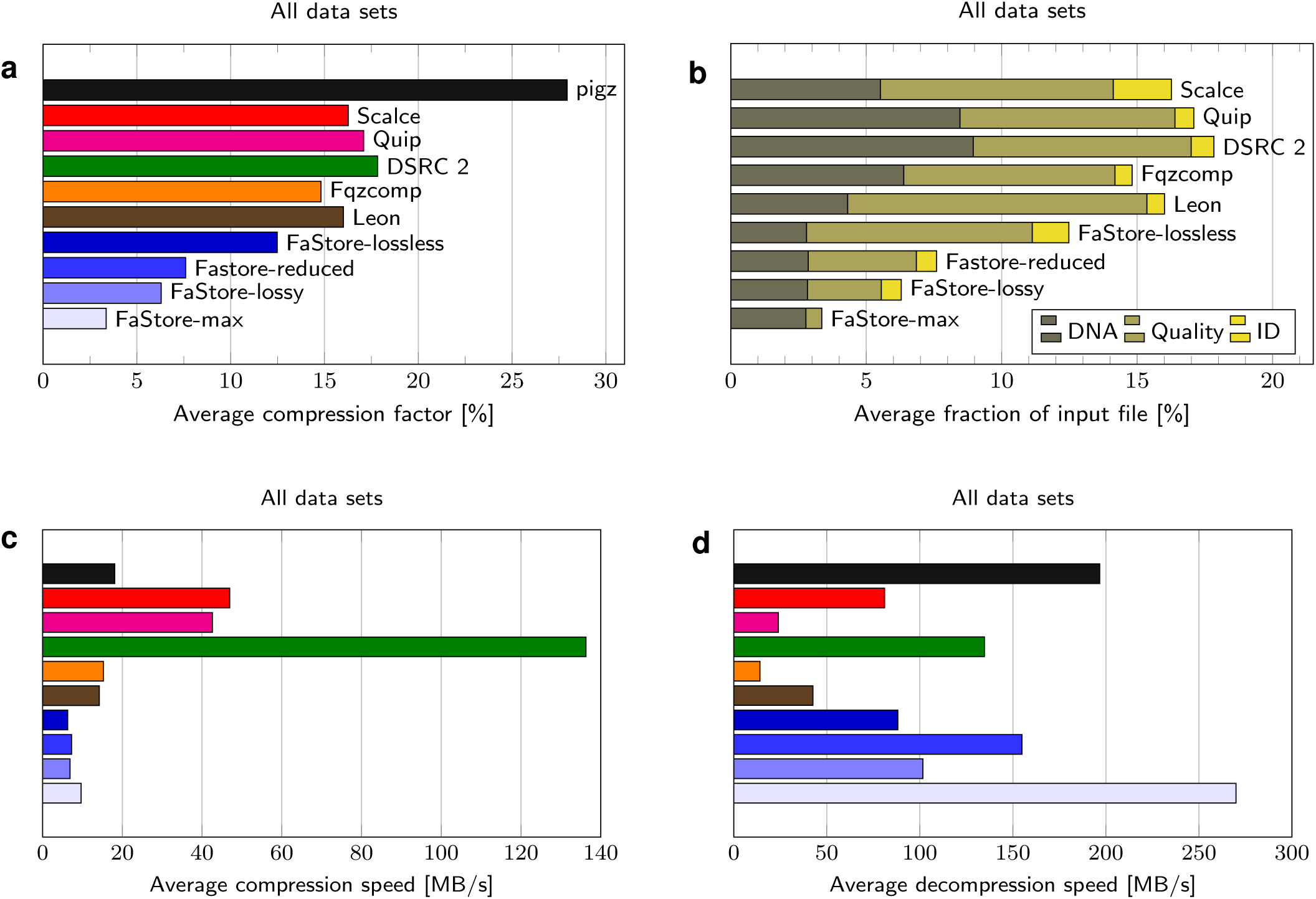
Compression results. (**a**) Average compression factors in % (compressed size divided by original size) for all examined datasets. (**b**) Average compression factors in % (compressed size divided by original size) for all examined datasets, divided by the different components: DNA bases, quality values, and IDs. (**c**) Average compression speeds for all examined datasets. (**d**) Average decompression speeds for all examined datasets. The sub-figures (**a**), (**c**), and (**d**) share the same legend. pigz is multithreaded variant of gzip (same compression ratios, but faster processing).

As shown in Figure 1a-d, FaStore in the *lossless* mode (preserving all the input data) achieves significantly better compression factors than the competitors. In particular, the compression gains with respect to the results achieved by the best competitor (i.e., Fqzcomp for all datasets except for dataset HSX where Leon outperforms Fqzcomp), range from 7.6% to 20.3%. For example, for *H.sapiens* datasets HS2 and WGS-42, this corresponds to more than 10GB of savings in both cases.

Although the *lossless* mode is used by default in FaStore, we strongly recommend considering some of the provided lossy modes. By discarding parts of the read identifiers and reducing the resolution of the quality scores, one can achieve significant savings in storage space. For example, in the *reduced* mode, the compressed size is about 60% of that of the *lossless* mode. The improvement is possible thanks to Illumina 8-level binning (Illumina 2014) and removal of the comments from the identifiers (which, in some cases, leads to storing only a library name and a read number). As the identifiers are usually truncated in this way by mappers when producing SAM files and the 8-level binning becomes a default option in modern sequencers (although the actual mapping of the values can depend on the internal configuration of the sequencing machine), this setting seems to be a reasonable choice.

Even better results (approximately half the size of the *lossless* mode) is possible when QVZ with distortion level 2 is applied (*lossy* mode). Nevertheless, the best compression factor (about a quarter of what was obtained in the *lossless* mode) is achievable when the identifiers are removed (only the pairing information between the reads is preserved) and the binary thresholding for the quality values is applied (*max* mode).

Figure 1b shows fractions of archives consumed by various components: DNA sequences, quality values, and read identifiers. There is no result for gzip as, due to the design of this compressor, it is impossible to measure the exact fraction of each component. The results show that FaStore uses much less space to store the DNA sequences than the competitors. Since in the lossy modes there is no loss incurred in the DNA sequences, the amounts of space necessary for storing them are almost identical across the different lossy modes. However, one needs to note that in the *lossless* mode FaStore needs more space for storing identifiers than the other competitors (except for Scalce, which also reorders the reads present in the FASTQ files to aid compression). The reason is that after reordering the reads it is much harder to compress their identifiers, as the neighboring ones differ more than in the original ordering. Nevertheless, the compression gain from the sequences of DNA overshadows the compression loss from the identifiers stream.

As outlined above, the most difficult to compress are, however, the quality scores. For most compressors, when the quality scores are losslessly compressed, they require more space than the DNA sequences and read identifiers together. Thus, applying the lossy schemes for the quality values has a remarkable impact on the total compression factor. For example, to losslessly compress *H.sapiens* dataset HS2, FaStore requires 46.3GB of space. From those, 32.7GB correspond to the quality scores, which can be further reduced to 9.3GB (Illumina binning, *reduced* mode), 8.4GB (QVZ with distortion level 2, *lossy* mode), or even 1.1GB (binary thresholding, *max* mode). Note that in all the cases the overall size of the lossless compressed FASTQ file is reduced by more than 50% when lossy compression of quality values is applied. In particular, a reduction from 46.3GB to as little as 14.7GB is achieved when binary thresholding is used. Furthermore, this reduction in total size is computed without considering lossy compression of the identifiers, which would provide even more storage savings. Below we demonstrate that such reductions in size are possible with little effect on variant calling.

Finally, Figures 1c–d show the compression and decompression speeds for the different methods analyzed in this paper. The compression speed is some drawback of our solution, as it is somewhat smaller than 10 MB/s (in the default C1 mode). Nevertheless, the decompression speed is comparable to the fastest algorithms, i.e., DSRC 2 and gzip. For use cases where compression speed is of uttermost importance, FaStore provides a fast mode, namely, the C0 mode. This mode trades the compression ratio of DNA sequences for compression speed, while still achieving better compression ratios than the competitors (see **Supplementary Worksheet W1**).

As reported in Table 2, when losslessly compressing FASTQ data, on average, the compression speed offered by C0 mode is greater by a factor of 5 as compared to C1 mode at a cost of increasing the size needed to store the DNA bases by a factor of 1.09. Note that switching between C0 and C1 has no significant effect on compression of quality scores and read identifiers, and the speed of decompressing files created in either of these modes is almost identical.

**Table 2:** Trade-off between modes C0 (fast) and C1 (default).

| Data set | Compression factor of DNA stream |  |  | Compression speed |  |  |
| --- | --- | --- | --- | --- | --- | --- |
|  | C0 [%] | C1 [%] | Ratio C0/C1 | C0 [MB/s] | C1 [MB/s] | Ratio C0/C1 |
| CE | 6.96 | 6.35 | 1.09 | 35.5 | 3.4 | 10.56 |
| GG | 8.96 | 8.41 | 1.07 | 39.2 | 6.2 | 6.30 |
| HS2 | 5.88 | 5.34 | 1.10 | 11.9 | 6.6 | 1.82 |
| HSX | 8.91 | 8.03 | 1.11 | 32.2 | 9.1 | 3.52 |
| WEX | 5.60 | 5.38 | 1.04 | 35.1 | 5.2 | 6.74 |
| WGS-14 | 10.82 | 10.28 | 1.05 | 32.5 | 5.1 | 6.39 |
| WGS-42 | 7.14 | 6.47 | 1.10 | 29.7 | 8.5 | 3.50 |

### Impact of lossy compression of quality scores on variant calling

Next, we assess the effect that the different lossy quality score compression modes provided by FaStore, namely, Illumina binning, binary thresholding, and QVZ, have on variant calling. Since QVZ optimizes the quantization for an average mean square error distortion level (specified as an input parameter), for the analysis, we considered distortion levels 1, 2, 4, 8, and 16.

For the evaluation, we selected the two datasets coming from whole-genome sequencing of *H.sapiens* individual, sequenced at coverage 14x (WGS-14) and 42x (WGS-42) (see Table 1). These datasets pertain to the same individual, namely NA12878, and were sequenced as part of the Illumina Platinum Genomes (Eberle et al. 2017). The reason for this choice is that the National Institute for Standards and Technology (NIST) has released a high-confidence set of variants for that individual as part of the Genome In a Bottle (GIAB) (Zook et al. 2014) initiative. This allows us to consider this set as the “ground truth” and use it to benchmark the different lossy modes supported by FaStore. For calling the variants we followed the Genome Analysis Toolkit (GATK) (McKenna et al. 2010) Best Practice pipeline (Auwera et al. 2013), utilizing BWA-MEM (Li and Durbin 2009) to map the reads to the human genome reference, followed by GATK HaplotypeCaller to call variants. We refer the reader to the **Supplementary Methods** for a detailed description of the pipeline used for the analysis.

In what follows we will report the results on Single Nucleotide Polymorphisms (SNPs), since SNPs are easier to detect and more curated in the high-confidence reference set. Nevertheless, for completeness, results for short insertions and deletions (INDELs) are provided in the **Supplementary Worksheet W2**. The GATK Best Practices proposes to apply VQSR for semi-automatic filtering of variants, however, the use of this machine-learning-based filter is still not widely adopted and should be used with caution for single-sample analyses. Hence, here we focus on the results obtained by applying hard filtering on the called set of SNPs, and refer the reader to the **Supplementary Worksheet W2** for the complete set of results achieved by applying both modes of filtering.

The results of the analysis are presented in Figures 2a–b. We focus on the recall vs precision results obtained when using the considered lossy modes for the WGS-14 and WGS-42 datasets, respectively. It is worth noticing that the precision is similar for both datasets (above 0.99 for most points), whereas the recall is much higher for WGS-42. The majority of points have a recall value around 0.981 for WGS-14 compared to 0.9978 for WGS-42. These results suggest that the sequence coverage plays a key role in variant calling as it improves with increasing coverage.

**Figure 2:**
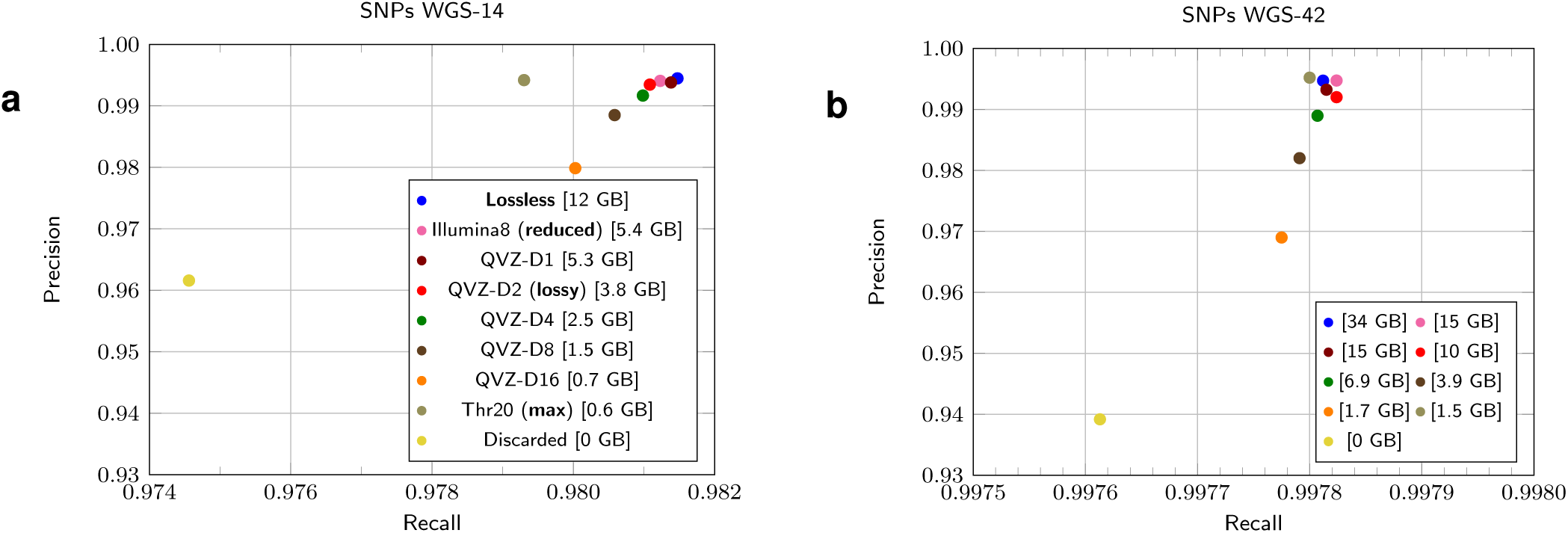
Compression results and variant calling analyses. (**a**) Results of variant calling for WGS-14 dataset. (**b**) Results of variant calling for WGS-42 dataset.

The most interesting aspects are, however, the results achieved using the various lossy modes. As expected, increasing significantly the distortion level of QVZ reduces both the recall and the precision. The variant calling performance applying QVZ with distortion level 1 (QVZ-D1) is comparable to that of Illumina binning, with slightly better results (in recall and in precision for WGS-42x) in favor of Illumina binning. Moreover, both modes, QVZ-D1 and Illumina binning, offer a similar compression factor, reducing the size of the quality scores by more than 44% when compared to lossless (with slightly better compression achieved by QVZ). In addition, the variant calling performance is almost indistinguishable from that achieved with the original data. Hence, the original size of the WGS-42 dataset could be reduced by an order of magnitude (a 12× reduction), and almost a half of the space required to store it losslessly compressed, by reducing the footprint of quality scores and removing the comments from the read identifiers, all while preserving the variant calling accuracy. As a side note, applying lossy compression to the read identifiers (as done in the *reduced* and *lossy* schemes) has no effect on variant calling, as the mappers usually strip the present comments, and can further reduce the compressed size.

In order to further boost the compression gains, more aggressive lossy modes need to be considered at the cost of possibly losing some variant calling accuracy. In that regard, QVZ applied with distortion level 2 (QVZ-D2) seems to be a good trade-off between variant calling performance and compression factor. It offers comparable performance to that obtained with the original data while reducing the size of the quality scores by more than 66%. Distortion levels above 2, although offer significant compression gains, show a degradation on variant calling. For example, with the maximum distortion considered (D16), the precision drops from above 0.99 (with the original data) to 0.98 (WGS-14) and 0.97 (WGS-42). The differences in recall are less pronounced.

Quite surprisingly, for WGS-42 the results for the *max* mode are almost as good as for the lossless mode (for WGS-14 the recall decreased by about 0.002), with a vast difference on the size of the compressed quality data (0.6GB vs. 12GB and 1.5GB vs. 34 GB, for WGS-14 and WGS-42, respectively). These results suggest that for data sets with increasing coverage, storing only the information of whether the called base is “good” or “bad” is sufficient for obtaining reliable results on variant calling, while significantly boosting the compression ratio.

For comparison, we also experimented with completely removing the quality data, but the results (series denoted as *discarded*) show a significant drop in both recall and precision. This indicates that some information about the base quality is necessary for reliable variant calling results (at least in the examined range of coverages).

### Impact of read reordering on variant calling

As mentioned above, the reads produced using next-generation sequencing protocols are randomly sampled from across the genome, and thus the original order of the reads carries no meaningful information. Due to the large size nature of the produced data, several commonly used computational methods that operate on these files rely on heuristics to be able to run in a reasonable time (even when executed in multi-threaded mode). For this reason, the reordering of reads, even if theoretically not relevant, may have some effect on variant calling. For example, authors in (Firtina et al. 2016) showed that, for some mappers, randomly shuffling the input FASTQ reads can lead to different alignment results, especially for reads originating from highly repetitive genomic regions.

Since FaStore permutes the input collection of reads, in this section we briefly examine the impact that various read reorderings can have on variant calling. The goal is thus to analyze how the FaStore-specific shuffling of the reads may affect variant calling. To that end, we compare it against a random shuffling of the reads in the file and against the original order. In addition, we included into the comparison the results from different computational environments (differing mainly by the number of computing threads employed). The framework for the evaluation is identical to the one employed in the previous section, that is, GATK pipeline is used as specified by the Best Practices using the data from *H. sapiens* NA12878 individual (see **Supplementary Methods** for a detailed description).

Figure 3 summarizes the findings of our study for dataset WGS-14. In particular, we compare the Precision and Recall (for the SNPs) with and without applying VQSR filtering, obtained with the original order, the FaStore order, and four random shuffles. For the original and FaStore orders, we run the experiments in two different computational environments (hence the two points with identical labels). There are several things to notice from this plot. First, the difference in precision / recall obtained by the various orderings is negligible. More interestingly, this difference is comparable to that of running the experiments in different computational environments, suggesting that the order has no more effect than the used computational environment. For example, when VQSR is not applied, the change in precision is less than 0.00006, and the change in recall is less than 0.002. On the other hand, applying VQSR increases the change in precision to 0.02, while maintaining the same variability in recall. More importantly, the choice of applying or not the VQSR filtering has significantly more effect (specially in Precision) than the ordering of the reads. The precision drops from 0.98 when no VQSR is applied to 0.91 when applied. This is several orders of magnitude larger than the change due to reordering of the reads. All this put together indicates that the gains in compression obtained by reordering the reads are worth pursuing, as there seems to be little – if none – effect on variant calling.

**Figure 3:**
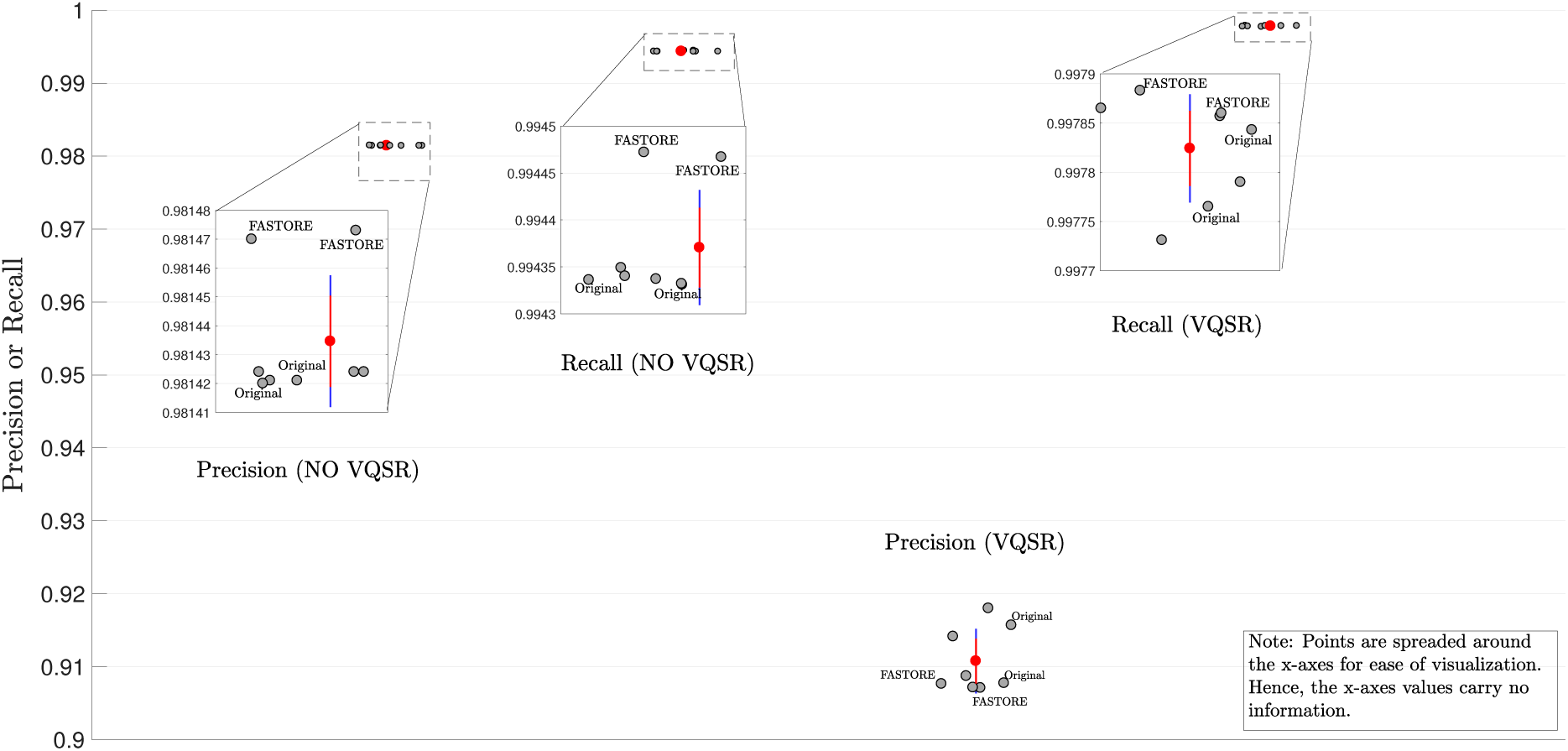
Precision and Recall obtained for various reorderings of dataset WGS-14, with and without VQSR filtering. Points without label correspond to random shuffling. The red point represents the mean, the red line is the 95% confidence on the mean, and the blue line is the standard deviation. The y-axis represents precision or recall, based on what it is specified in the x-axis.

### Influence of coverage

Finally, we analyzed the compression ratio just for the DNA symbols (Figure 4) using whole-genome sequencing data of *H.sapiens* (WGS-235 dataset) sampled at various coverages. As can be noted, for some algorithms (Scalce, Leon, Fqzcomp, FaStore) the increasing coverage leads to significant improvements in compression ratio. In the case of FaStore, the advantage is more than 2-fold over the competitors. When testing read-reordering algorithms (FaStore and Scalce), we also added a series of values in which the reads were compressed as single-end (i.e., the pairing information was lost). For FaStore this led to further savings in storage space, obtaining about 2.6 times better compression ratios.

**Figure 4:**
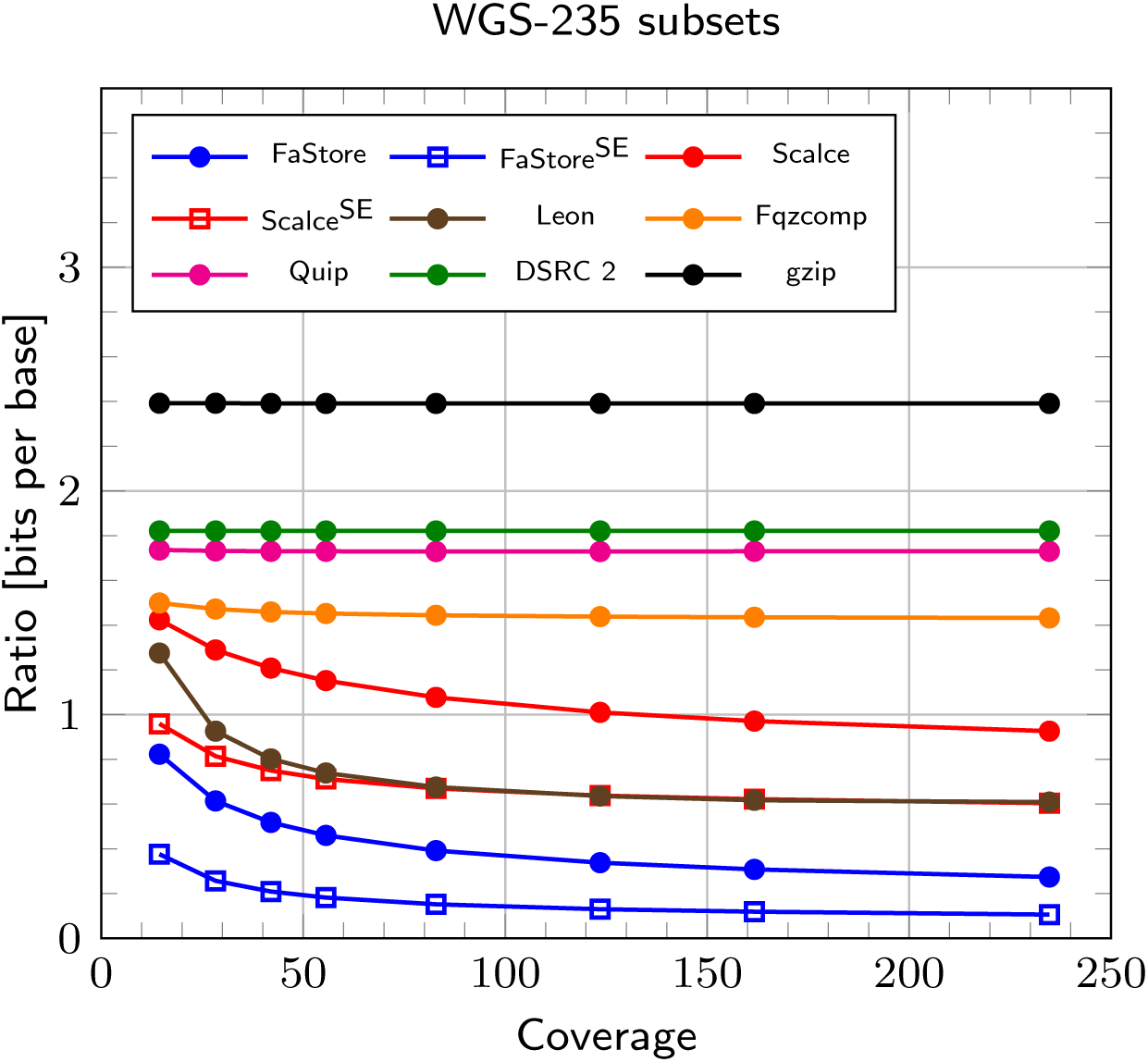
Compression ratio for storing only DNA symbols (bits used to encode a single base) for *H.sapiens* sampled at various coverages (WGS-235 subsets). Superscript SE stands for Single-End.

## Discussion

The efficient storage and transfer of huge files containing raw sequencing data has become a real challenge. The popular general-purpose compressor gzip is still being used as a *de facto* standard solution to compress FASTQ files, being able to reduce file sizes by about 3 times, with significant gains in cost of storage and speed of transfer. However, as has been already pointed by multiple researchers (Numanagić et al. 2016; Roguski and Ribeca 2016), even just by splitting the content of FASTQ file into separate information streams of homogeneous data type (i.e., DNA sequences, read identifiers, and quality scores) and compressing each separately using gzip, one can achieve gains in compression more than 15% than by using gzip alone. Unfortunately, this clearly demonstrates that the current *de facto* standard approach for storing raw sequencing data is not the best choice and in modern times much higher savings in storage are possible and necessary.

Therefore, our proposed compressor, FaStore, is designed to achieve excellent compression factors, i.e., about 3 times better than gzip and significantly better than the existing specialized FASTQ compressors. In addition, FaStore offers several lossy compression modes for the quality scores and the read identifiers, which result in significant compression gains. Moreover, since the methods to compress DNA sequences, quality scores and read identifiers are modular, they can be possibly implemented in other solutions to store sequencing data in a more compact form. For example, the DNA sequences compression methods can be used in solutions working with SAM alignments to improve the compression of unaligned reads. The methods to compress quality scores can also be applied when compressing alignments in SAM format. Finally, all the methods can be used in different genomic data processing and compression frameworks, e.g., implemented as codecs in CARGO (Roguski and Ribeca 2016) or Goby framework (Campagne et al. 2013).

In parallel, we strongly suggest the community to consider resignation from storage of all the raw sequenced data or, at least, considering the raw data to be stored in one of the lossy forms. As we presented, together with the increasing sequencing throughput and the dropping costs of sequencing reflected in higher coverages for the smaller prices, the high resolution of quality values seems to be unnecessary. An important stage in this direction was made by Illumina, which has already introduced the possibility of reducing the resolution of the available quality scores to 8 values in some of their newest sequencers (e.g., HiSeq X), and it is considering a more aggressive 4-level binning scheme for their latest NovaSeq system. We show that similar variant calling results could be obtained when even more reduction of the quality stream is applied. For sufficiently large coverage it seems to be enough to provide just a binary information about each base telling whether it is “good” or “bad”.

To imagine the possible gains in reduction of cost thanks to the lossy approaches let us say that the FASTQ files for *H.sapiens* sequenced at 42-fold coverage in the paired-end mode could consume as little as 10 Gigabytes (FaStore-*max*), which can be compared to 110 Gigabytes of gzipped FASTQ files. For both datasets the quality of variant calling results should be almost identical.

## Methods

### Compression workflow

In FaStore, the compression workflow has been designed as a multi-step process to exploit the high sequence redundancy present in the sequencing data. It consists of: (1) reads clustering, (2) optional reads re-clustering, and (3) reads compression stages. Each stage is further divided into multiple smaller steps. In this section we provide a general overview of the compression workflow. A detailed description of the methods used to cluster the reads and to compress the DNA sequences, quality scores, and read identifiers can be found in the **Supplementary Methods**. The workflow is depicted in Figure 5 and can be briefly described as follows.

**Figure 5:**
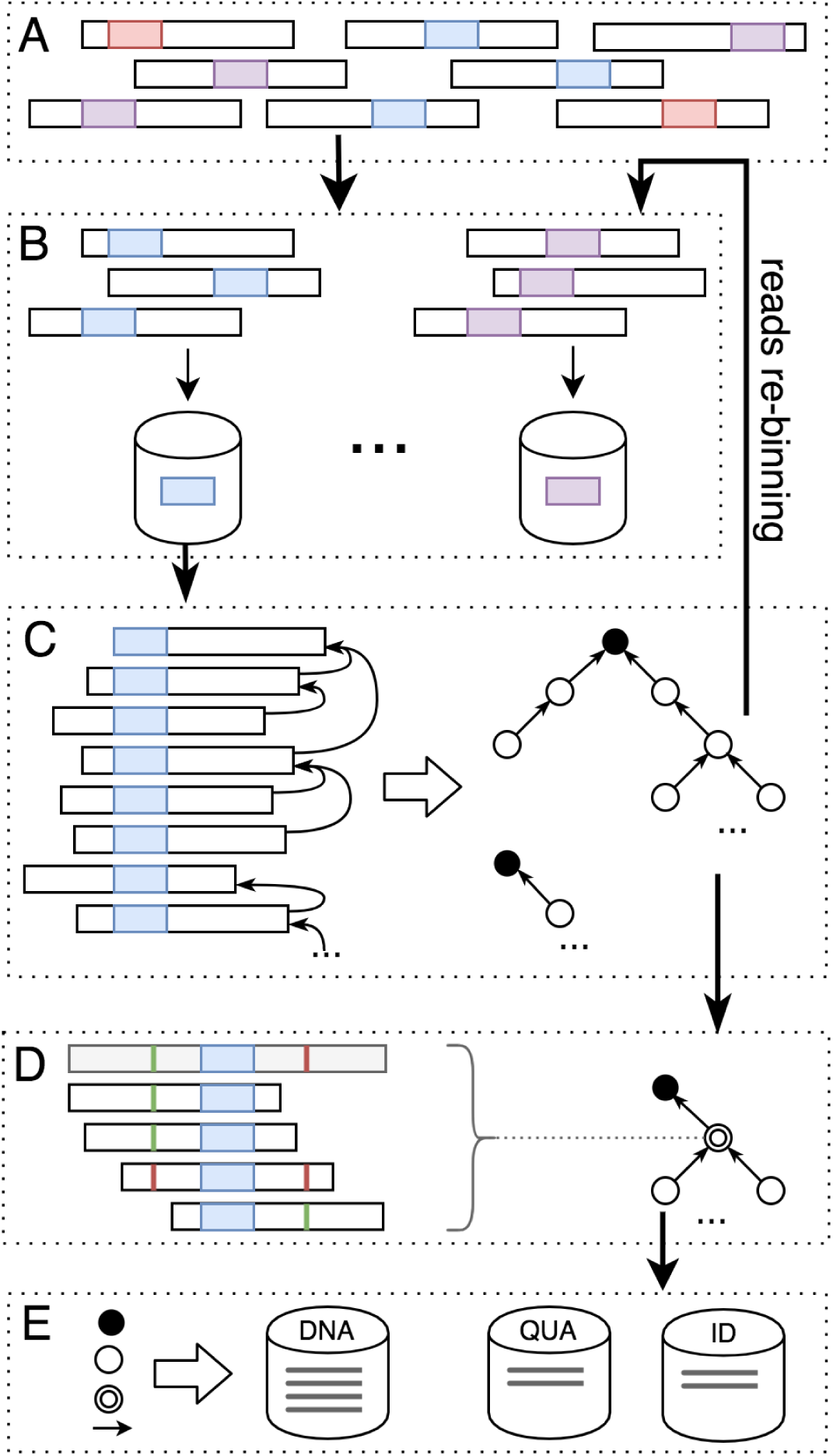
General compression workflow of FaStore. (**A**) Raw FASTQ reads are (**B**) distributed into bins. (**C**) Within each bin, the reads are matched, giving as a result a reads similarity graph. Optionally, the reads follow further redistribution and matching. (**D**) With the final similarity graph, the reads are assembled into contigs. (**E**) The reads are encoded either in contigs or differentially, depending on the matching result.

The read clustering stage is a 2-step process, consisting of a read binning step (see Figure 5B) and a read matching step (see Figure 5C). During binning, for each read from the input FASTQ file(s) (Figure 5A), FaStore seeks the sequence signature, i.e., the lexicographically smallest *k*-mer, with some restrictions. The signature is used as an identifier of the bin in which the read is placed into. During this step, some statistics are gathered related to the observed DNA sequences, quality scores, and read identifiers. At the end of the binning stage, these statistics are used to compute the quantizers for the quality scores, which are stored in a global codebook. This process is performed only when compressing quality scores in QVZ mode. In addition, a token dictionary is built for the read identifiers. These data will be used during the final compression stage.

After binning all the reads, FaStore performs the matching process, independently per each bin. The goal is to find for each read, a referential one, which has the lowest “encoding cost”. This cost corresponds to the number of operations required to transform one sequence into another under some user-specified constraints. In order to do so, FaStore first reorders the reads, so that reads with DNA sequences possibly originating from the same genomic region, are likely placed close to each other. Then, we iterate over the reordered reads and, for each sequence, we search in a window of *m* previous ones for the best match. A read can be matched as a *normal match*, an *exact match* (an identical sequence was found), or as a *hard read* (when no satisfactory reference was found). The result of reads matching is represented as a similarity graph, where each node represents a read (DNA sequence) and the edge represents the type of match. More specifically, the result is a collection of trees, where each hard read represents a tree root (a tree can also consist of only a root node). With such graph we can already proceed to the compression stage (as in C0 mode).

In order to improve the clustering between the sequences, a number of optional reads re-clustering steps can be performed (C1 mode). The goal is to create larger clusters of highly similar (groups of) reads to possibly bring the reads from the same genomic regions close to each other, by re-distributing the reads. To do so, we first define a new subset of signatures, which will be used as a filter, to select the bins into which the reads can be moved. Then, for each tree, we select a new root node, which has a new signature residing at the beginning or at the end of its sequence. The connections between nodes are updated and the trees are moved into bins (similarly as in Figure 5B) denoted by their root signatures, where each tree is represented in the new bin as a single read (its root). This allows to improve the clustering between the reads, by performing an additional matching of them (as in Figure 5C) and, as a result, building larger trees of similar reads. In the C1 mode three re-distribution step(s) are performed.

The compression stage is a two-step process, consisting of assembling the reads sequences (Figure 5D) into contigs and encoding the reads (Figure 5E), using the previously built reads similarity graph. First, we traverse each tree and try to assemble the reads into possibly large contigs. The goal is to encode the reads with respect to the built consensus sequences, encoding only the variants (if present) in the contigs. While assembling a contig, for each read, we try to anchor it into the consensus sequence using the position of its signature (which resides at the “center” of the consensus). To add the read to the contig, we assess whether it does not introduce too many variants into the current consensus sequences, as they will need to be encoded by the other reads already present in the contig. When no more reads can be added to the contig, its final consensus sequence is determined by majority voting. As a result, in the graph some of the nodes are replaced with the contig nodes, updating the connections between nodes accordingly.

Finally, we proceed to encode the reads data (Figure 5E), storing the result in a number of streams, separately for DNA sequences, quality scores, and read identifiers. The read sequences are encoded either in contigs (encoding differentially versus consensus sequences) or differentially versus each other, depending on the matching result. To encode the quality scores using QVZ we use the quantizers from the previously created codebook. Alternatively, when using Illumina 8-level binning or binary thresholding, we encode the transformed quality values. In parallel, we encode the read identifiers using the previously built dictionary. Finally, the streams are compressed using a custom arithmetic coder or the general-purpose compressor PPMd.

### Variant calling

To investigate the possible side effects of applying lossy compression for base quality scores, we first prepared a set of test FASTQ files, WGS-14 and WGS-42, which come from deep sequencing of the NA12878 *H.sapiens* individual. These files included, in addition to the original input files: (a) original input files (lossless), (b) original input files with lossy compressed quality scores, (c) FaStore-shuffled reads with lossy compressed quality scores. Moreover, using WGS-14 dataset we tested the effect of reordering the reads using an additional set of test FASTQ files (but without applying any compression). These included: (d) original input files with randomly shuffled reads, (e) FaStore-shuffled reads. A detailed description of the FASTQ files preparation steps can be found in **Supplementary Methods**.

With such prepared input FASTQ files, we followed the GATK (McKenna et al. 2010) Best Practices recommendations (Auwera et al. 2013) to assess the variants. We used BWA-MEM (Li and Durbin 2009; Li 2013) to map the reads to the human genome assembly GRCh37. Following a number of post-processing steps, we called the variants using GATK HaplotypeCaller (GATK-HC). For assessing the variant calling performance, we used as a “gold standard” the variants for NA12878 provided by the GIAB (Zook et al. 2014), and benchmarked our results using the Illumina Haplotype comparison tools pipeline (https://github.com/Illumina/hap.py). This pipeline is also recommended by the Global Alliance for Genomics and Health (GA4GH) as one of the benchmarking standards. In the manuscript we reported precision and recall results for the obtained SNPs. For completeness, we also filtered the variants using GATK Variant Quality Scores Recalibration (VQSR). Both the SNPs and INDELs calling results, with and without VQSR filtering, are available in **Supplementary Worksheet W2**.

## Software availability

FaStore can be downloaded from https://github.com/refresh-bio/FaStore.

## Acknowledgments

We would like to thank Ivo Gut for supporting the project and Marcos Fernández-Callejo for helpful discussions and technical insights.

This work was supported by: National Science Centre, Poland [under project DEC-2016/21/B/ST6/02153 to S.D.]; European Unions Seventh Framework Programme (FP7/2007-2013) [under grant agreement No. 305444 (RD-Connect) to Ł.R.]. The infrastructure was supported by “PL-LAB2020” project, contract POIG.02.03.01-00-104/13-00.

## Competing financial interests

The authors declare no competing financial interests.

